# The use of phylogenetics and ancestral sequence reconstruction to identify improved halophilic enzymes for the biodegradation of poly(*R*-3-hydroxybutyrate)

**DOI:** 10.1101/2022.10.12.511935

**Authors:** Stephen Quirk, Kassi Kosnicki

## Abstract

Evolutionary analysis combined with ancestral sequence reconstruction (ASR) was utilized to calculate the taxonomic relationship between poly(hydroxybutyrate) depolymerase (PHBase) enzymes belonging to the halophilic genus *Halomonas*. Enzymes cluster into four distinct clades that differ in sequence length and composition. Like many of the previously studied PHBase proteins, there is a high degree of heterogeneity at the genus level. Ancestor sequences were calculated for each tree node using a maximum likelihood approach. The most ancestral sequence and one extant enzyme from each of the clades were expressed in *Escherichia coli*, purified to homogeneity, and characterized. The ancestral enzyme binds substrate more efficiently, is kinetically faster, and is more stable at elevated temperatures compared to the extant enzymes. Hence, an evolutionary phylogenetic approach is a viable alternative to other techniques for identifying optimized enzymes for industrial PHB degradation.

## 1. Introduction

The use of petroleum-based, non-degradable plastics, like polypropylene and polyethylene has become a target for social activism and growing government regulation. As ocean accumulation increases and available land for landfills decreases, the continued reliance on this class of polymer will no longer be an option in the global production of consumer plastics. As the worldwide single-use plastic epidemic worsens, it becomes paramount to identify fully renewable plastics. Much work has been accomplished on the family of polyhydroxyalkanoates [1], most notably polyhydroxybutyrate (PHB; for a review see [2]). The polyhydroxyalkanoates represent viable replacements for polypropylene and polyethylene, as they are fully bio-renewable, biocompatible [3], and are produced across many bacterial, fungal, and archaeal lineages [3]. PHB also has processing characteristics that most closely mirror those of polypropylene, yet with some notable differences in flow viscosity[4]. Regardless of their source, PHBs are synthesized in a three step enzymatic process beginning with acetyl-CoA. The first step is catalyzed by PhaA (Acetyl-CoA acetyltransferase; EC 2.3.1.9) which forms acetoacetyl-CoA. This in turn is converted into R-3-hydroxybutyryl-CoA by the PhaB enzyme (Acetyl-CoA reductase; EC 1.1.1.36) in a NADP-dependent reaction. The final step, catalyzed by PhaC (PHB polymerase; EC 2.3.1.-), is the polymerization of R-3-hydroxybutyryl-CoA into PHB (see [5]). PHB biodegradation is accomplished by an extra- or intracellular PHB depolymerase, PhaZ [6]. The biodegradation of PHB results in the formation of the monomer hydroxbutyric acid and small PHB oligomers [7] according to Figure 1. The reaction is catalyzed by a single enzyme.

**Figure 1.**
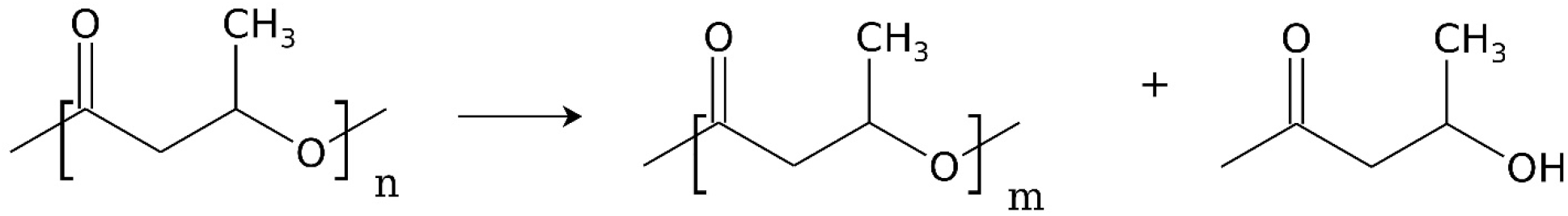
Enzymatic degradation of poly(*R*-3-hydroxybutyrate) by PHB depolymerase. For this scheme m<<<n and where n represents small oligomers.

PHB depolymerase (E.C. 3.1.1.75), is a carboxylic ester hydrolase. Although the enzymes have been cloned and studied from a variety of organisms, including both mesophilic and thermophilic bacteria from a wide variety of environment (*e.g*.- [8–13]) there are issues with the enzymes – primarily the large number of disulfide bonds and the observation that they are kinetically slow. In addition, the scale-up and purification of PHB granules from both meso- and thermophiles has been problematic under aseptic conditions. For a general review of known PHBases see [14].

The genus *Halomonas* constitutes a group of halophilic (salt tolerant) bacterial species that are capable of producing PHB as a potential energy sink, as well as to help coordinate life in an environment characterized by high osmotic stress [15], and other stressors [16]. To date there are 79 species within the genus with varying degrees of salt tolerance, typically in the range of 5 – 25% (w/v) NaCl, and differing metabolic phenotypes [17]. *Halomonas* species are prime candidates for sourcing PHB as granule purification is simplified by employing a hypotonic cell lysis regime [18] and as such can play an important role in the production of many chemicals, proteins, and other compounds. In fact, *H. bluephagenesis* is emerging as a promising candidate for synthetic biology approaches to customize bioproduction [19]. Although several reports discuss the kinetics of PHB formation in several *Halomonas* species including *H. halophila* [20], *H. elongata* [21], *H. boliviensis* [22], TDO1 [23], KM-1 [24], and *H. nitroreducens* [25] there are no reports that characterize PHB biodegradation in this genus. Since it is advantageous to run bioreactor processes in high salt in lieu of other aseptic techniques or sterilization, it is important to characterize the other PHB pathway enzymes from *Halomonas* as a first step in creating a bioreactor process that can not only generate PHB, but degrade it, and reform it. That is a truly circular process.

The traditional route to fine-tuning the kinetic and stability parameters of an enzyme is through the use of site directed mutagenesis, often employing a high throughput methodology [26] or by directed evolution methods [27]. An emerging method that can be used separately or in conjunction with these techniques is through the reconstruction of ancestral sequences [28] [29] [30] [31]. Ancestral sequence reconstruction (ASR) has successfully identified enzymes with increased thermostability (*e.g.-* [32], increased substrate binding [33], altered substrate specificity [34], protein activation [35],as well as alterations to subunit composition in enzyme complexes [36]. ASR can also be utilized as a high throughput method [37], has been used to modify protein drugs [38]and is useful when used in conjunction with protein design [39]. The method has also been used to design *de novo* active sites into ancestral protein backbones [40] and can shed light into folding mechanisms [41,42]. Importantly, ASR has been shown to produce enzymes with functions closer to their extant counterparts than the use of consensus sequences derived through multiple sequence alignments [43].

This work seeks to determine if evolutionary analysis coupled with ancestral sequence reconstruction can identify halophilic PHBases with improved kinetic efficiency and increased stability versus their extant counterparts. This would serve as a counterpart to site directed mutagenesis or directed evolution methods to optimize/identify an enzyme better suited for bioreactor-based biodegradation of PHB. The work shows that starting from a single genus of interest it is indeed possible to perform a phylogenetic analysis of the sequences in order to better understand the evolutionary history of this enzyme and the resulting sequence variation over time within the genus *Halomonas*. Analysis of the ancestral sequences at each node in the evolutionary tree identifies multiple possible alternatives to the extant sequence. Production of one such ancestral sequence results in an enzyme that can degrade PHB faster (as measured both in the binding and in the chemical step) and is more stable.

## 2. Materials and Methods

### 2.1 Materials and chemicals

All chemicals were purchased from Sigma-Aldrich (St. Louis, Mo USA), including all buffers, media components, isopropyl-β-D-thiogalactoside, antibiotics, and polyhydroxybutyrate. Chromatography resins were from Sigma-Aldrich or BioRad, Inc. All laboratory supplies were purchased from Fisher Scientific. Competent *E. coli* were purchased from New England Biolabs, Inc. Carboxylate-modified polystyrene particles we obtained from Creative Diagnostics. Nickel-NTA modified spheres were obtained from Polysciences, Inc.

### 2.2 Sequence collection and phylogenetic analysis

The NCBI Identical Protein Groups database was queried with the search string “PHB and Depolymerase and Halomonas”, with a sequence cutoff range of 300 – 1000 amino acids to avoid fragments. The sequences were downloaded into a single FASTA formatted file. GUIDANCE2 [44] was utilized to identify any sequences that were negatively contributing to the quality of the multiple sequence alignment (MSA) using either PRANK [45] or MAFFT [46] alignments in order to optimize the quality of the MSA [47]. A sequence cutoff value of 0.7 was applied. A final multiple sequence alignment (MSA) file was prepared using MUSCLE [48]. A maximum likelihood (ML) evolutionary analysis was performed using RAxML [49,50] with a JTT substitution matrix, ML estimated stationary base frequencies, and a GAMMA among-site rate heterogeneity model with mean category rates. During the ML search tree topology, branch lengths, and tree models were optimized. A consensus tree was calculated from 1000 bootstrapped replicates. Ancestral sequences were reconstructed at each node in the tree using the program ProtPars [51]. Some additional ASR analysis was performed with MEGA [52]. Tree diagrams were created with FigTree (http://tree.bio.ed.ac.uk/software/figtree/) and analyzed using TREE2FASTA [53].

### 2.3 Protein construct design

Standard molecular biology PCR techniques were employed to produce expression constructs for the enzymes chosen from the evolutionary analysis. The amino acid sequences were reverse translated and codon optimized for expression in *E. coli*. Solubility and folding were assisted by cloning the open reading frames downstream of a gene encoding maltose binding protein (MBP). A hexahistidine purification tag was placed at the 5’ start of each open reading frame and a TEV protease cleavage sequence was placed between the MBP and PHBase sequences. Hence each expression construct was in the form of [His6-MBP-TEV-PHBase]. The gene inserts were cloned into an expression vector (amp^r^, medium strength ribosomal binding site). The inserts were verified by DNA sequencing after construction.

### 2.4 Enzyme purification

The expression plasmid was used to transform chemically competent Origami-2(DE3) bacteria. Single colonies were selected from LB-Amp plates and used for expression screening. Colonies were grown at 37 °C for 12 hours in LB media supplemented with 100 μg/mL ampicillin. This culture was used to inoculate fresh LB-Amp flasks at a 1:100 inoculum. These cultures were grown at 25 °C until OD_595_ = 0.4 (typically 4 hours) at which time IPTG was added to a final concentration of 1 mM. Growth was continued for 16 hours. Cells were harvested by centrifugation at 10,000 xg for 15 minutes and frozen at −80 °C until use (minimal time frozen was 24 hours). Cells were thawed on ice and were resuspended in buffer: 0.5 M NaCl, 20 mM Tris-HCl, 5 mM imidazole, pH 7.9 (typically 1 mL per gram of cells). Cells were disrupted via two passes through a French Press followed by centrifugation at 30,000 x g for 30 minutes. The supernatant was slowly passed over a 5 cm x 4.9 cc His-Bind resin column (bead height 3 – 4 cm). The column was washed with 10 column volumes of wash buffer (0.5 M NaCl, 20 mM Tris-HCl, 60 mM imidazole, pH 7.9) at a flow rate of 0.4 mL/min. PHBase was eluted from the column with the addition of 3 column volumes of 0.5 M NaCl, 20 mM Tris-HCl, 1.0 M imidazole, pH 7.9. The pooled fractions were applied to a 70 cm x 4.9 cc Sephadex G-100 column (10 mM Tris-HCl, pH 7.5, 1 mM EDTA). Fractions containing homogeneous PHBase were pooled (after inspection by SDS PAGE), concentrated to 10 mg/mL via Centricon filters. Enzyme was stored frozen at −20 °C until use. The histidine tag/MBP fusion was removed from the enzyme using TEV protease. Protein was diluted to 1.0 mg/mL into 10 mM Tris-HCl, pH 7.5, 25 mM NaCl. 100 U of TEV protease was added per mg of enzyme (approximate ratio of 1:100 (w/w). The reaction was allowed to proceed for 16 h at 4 °C. The mixture was passed over a charged nickel column. One column volume of flow through was collected as purified tag-free enzyme.

### 2.5 Enzymatic reaction conditions

A turbidometric assay was employed to measure PHBase activity under various conditions. The standard reaction (final volume = 1.0 mL) contained 200 mg/L of PHB granules (that were previously stably suspended via sonication), 10 mM CaCl_2_, 25 mM buffer at various pH values, 250 mM NaCl, and 100 mM KCl The reaction was initiated after the addition of enzyme and monitored at 650 nm in Applied Photophysics spectrapolarometer in absorbance mode. The reaction was gently stirred and maintained at a constant temperature. OD measurements (typically starting in the range of 3-4) were converted to percent OD remaining as a function of time. Alternatively, a second assay was utilized to measure β-hydroxybutyrate directly using the Sigma-Aldrich hydroxybutyrate assay kit MAK272. HB was measured fluorometrically (λex = 535 nm, λem = 587 nm). Aliquots (10 μL) were removed from the PHB depolymerase reaction at various time points, mixed with 50 μL of the supplied HB assay buffer, and pipeted into a well of a black, flat bottomed, 96-well plate. The plate was incubated at room temperature in the dark for 30 minutes. Fluorescence emission intensity was measured using a Molecular Dynamics SpectraMax M5. Fluorescence readings were converted to HB concentration via comparison to a standard curve constructed from known concentrations of pure hydroxybutyrate. All kinetic parameters are calculated per [54].

### 2.6 Tethering the enzyme to a surface

PHBase (as intact fusion protein) was immobilized to the surface of a high capacity nickel nitrilotriacetate polystyrene bead (10 μm diameter) by incubating the sphere in a solution of PHBase (100 μg/mL) in 0.3 M NaCl, 20 mM Tris-HCl, pH 7.5 at 4 °C for 12 h. The beads were briefly rinsed in buffer devoid of enzyme and then placed into the reaction tube. At no time were the beads exposed to air for more than 1-2 secs. Once tethered the enzyme coated beads were placed into a 20 mL vial, on a screen support, above a stir bar, immersed in reaction buffer with a 1.0 cm^2^ piece of PHB film. The reaction vial was placed into an incubator on a stir plate and the reaction was allowed to proceed at 40 °C. Aliquots were taken out as a function of time and measured for fluorescence at 587 nm, indicating the amount of HB liberated from the PHB film.

## 3. Results

### 3.1 Evolutionary analysis of the genus Halomonas

The NCBI database search returned 82 unique PHBase sequences (all putative enzymes). GUIDANCE2 analysis suggested rejecting three sequences due to poor fit statistics (at seq_cutoff values of 0.7) using the PRANK alignment algorithm. The remaining 79 sequences constituted the final database. A Python script was used to determine *Halomonas* PHBase population statistics. The mean sequence length (and standard deviation) was 368 +/− 54 amino acids with an isoelectric point distribution of 5.7 +/− 1.0. The 79 sequences contained on average 11.6 +/− 1.6 percent negative amino acids, 8.2 +/− 1.0 percent positive amino acids, and 7.5 +/− 0.7 percent aromatic amino acids. The distributions for these descriptors are shown in Table I for each clade as well as the entire genus. Overall there is a fair degree of sequence length and composition heterogeneity in the genus, as is seen when looking at the sequences of other published PHBases. Sequence identity across the genus ranged from 48% to 97%.

**Table 1.** Sequence properties of *Halomonas* taxonomic groups.

| Clade | Amino Acids |  | pI |  | Cysteines |  |
| --- | --- | --- | --- | --- | --- | --- |
|  | Ave. (SD) | Range | Ave. (SD) | Range | Ave. (SD) | Range |
| All | 368 (54) | 300-494 | 5.7 (1.1) | 4.2-8.5 | 7.5 (0.7) | 0-11 |
| I | 396 (14) | 374-411 | 6.5 (1.0) | 5.4-8.5 | 5.0 (1.0) | 4-7 |
| II | 440 (17) | 395-454 | 6.5 (0.7) | 5.8-8.4 | 5.2 (1.0) | 4-7 |
| III | 348 (16) | 300-368 | 4.6 (0.2) | 4.2-4.8 | 8.3 (1.0) | 8-10 |
| IV | 302 (2) | 300-306 | 5.1 (0.2) | 4.8-5.7 | 1.7 (0.6) | 1-3 |

The ML consensus evolutionary tree is presented in Figure 2 and is calculated from 1000 bootstrap replicates. The percentage of tree replicates characterized by a particular branch is noted on a tree in the Supplemental Data file. Branch replicates range from 18 to 100 percent. The consensus tree is characterized by 75 nodes, of which 76% are seen in at least 50% of the replicate trees. The consensus tree can be subdivided, based on branching, into four primary groups or clades. Each of which is characterized by different sequence descriptors as is seen in Table I. This analysis is not a strict evolutionary history as the tree is unrooted, yet it is useful to identify ancestral sequences at various nodes of interest. The divergence time across the entire genus was 3.1 million years as estimated using the timed tree function in MEGA. For each clade, one extant sequence was chosen as a representative of that clade (marked with a highlighted boxed accession number in Figure 2). The sequence that was closest to the clade sequence descriptor average (length, pI, charged residues, number of cysteines) was selected. All ancestral sequences were visually inspected and the most ancestral sequence (denoted by the black circle in Figure 2) was chosen for further study. The extant and ancestral sequences are identified in Table II. An alignment of the four extant clade enzymes and the ancestral sequence (ANC1) is shown in the supplementary Data file. ANC1 ranges from 21.1% identity (eIII) to 85.9% identity (eI) as is shown in Table II.

**Figure 2.**
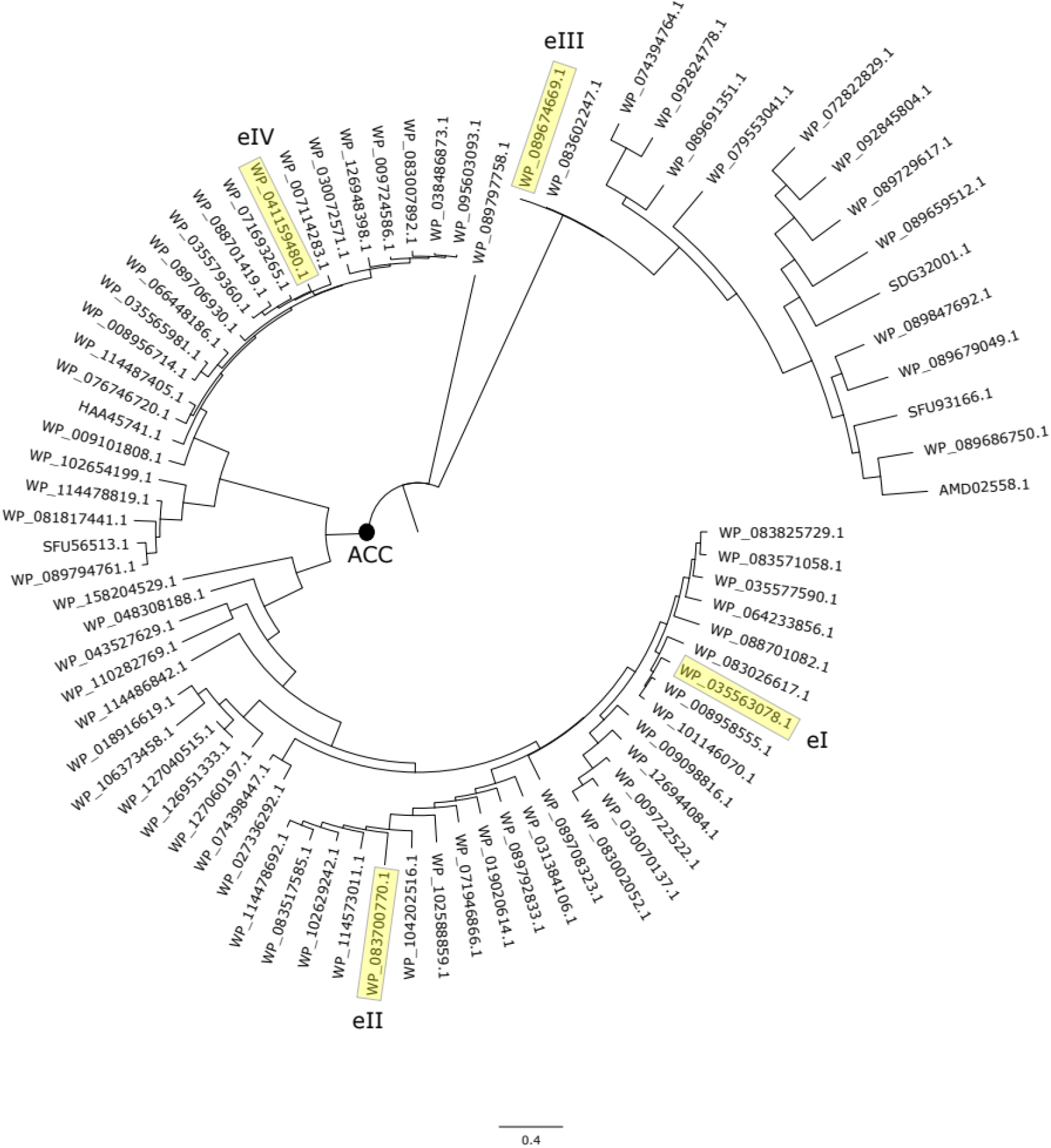
Unrooted Maximum Likelihood consensus tree. 1000 bootstrapped replicates. Four clades (I – IV) with extant (e) representative are highlighted in yellow. The ASR sequence derived from the most ancestral node is identified with a black circle.

**Table 2.** Identification of *Halomonas* PHBase sequences

| Extant Clade | Accession Number | Species | pI | Mol. Weight<br>(kDa) | %ID<sub>ANC1</sub> |
| --- | --- | --- | --- | --- | --- |
| eI | WP_035563078 | K0116 | 5.41 | 43.4 | 85.9 |
| eII | WP_083700770 | 1513 | 5.96 | 44.4 | 74.5 |
| eIII | WP_089674669 | aquamarina | 4.54 | 36.1 | 21.1 |
| eIV | WP_041159480 | KHS3 | 5.63 | 32.6 | 31.8 |
| ANC1 | - | - | 6.21 | 44.3 | - |

### 3.2 Cloning and expression of PHBase enzymes

The protein over expression vectors efficiently produce the four extant and one ancestral enzymes when induced by IPTG. Low temperature induction resulted in better yield than growth at temperatures closer to 37 °C (data not shown). Typical yields are as follows: eI-10.2 mg/L, eII-14.6 mg/L, eIII-11.8 mg/L, eIV-16.6 mg/L, and ANC1, 22.5 mg/L; where ‘e’ refers to the extant enzyme from clades I – IV, and ANC1 is the ancestor reconstructed enzyme. Presumably the slower induction at lower temperatures allows for better protein folding and lower aggregation. The Origami-2(DE3) expression host was critical in allowing disulfide formation as no soluble protein was produced in a regular BL21(DE3) strain (data not shown). No further experiments were performed to optimize expression levels. The hexahistadine tag greatly facilitated purification to a two step process which resulted in homogeneous enzyme. The entire affinity tag was also efficiently removed with TEV protease. All enzyme assays were performed on cleaved PHBase. Figure 3A shows a gel electrophoresis analysis of the final purified enzymes used in this study. Of note, and for unknown reasons, the eII protein aberrantly migrates on the gel as a smaller sized protein. The primary sequence of the reconstructed ANC1 enzyme is shown in Figure 3B. The sequence contains the four PHBase cannonical consensus structures that surround active site residues. Structurally, the ANC1 enzyme can be modeled by threading the ANC1 sequence onto the backbone of a known homolog, which returns a best fit 3-D all atom model. The selected PDB structure for the thread was 2D80, which is the PHBase from *Penicillium funiculosum*. Once threaded the model was subjected to 5 ns of NVT molecular dynamics (in solvent) to relax the model. The final ANC1 model has an Rmsd of 8.7 A for overlapping mainchain atoms compared to 2D80. ANC1 is primarily characterized by the central parallel beta sheet, which is a hallmark of known PHBase structures. The model is shown in Figure 4A and the superposition with 2D80 is shown in Figure 4B.

**Figure 3.**
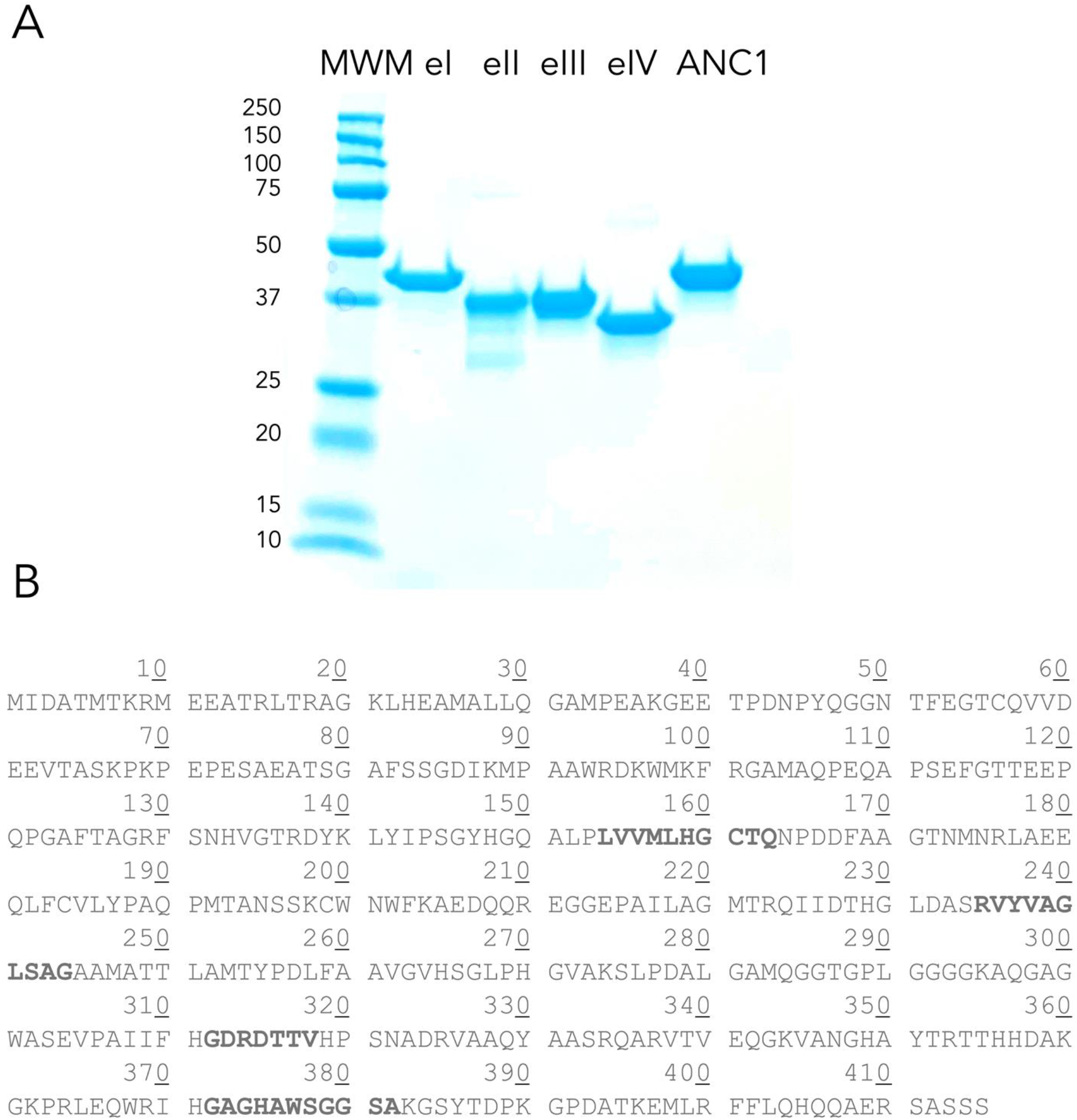
**A)** Gel electrophoresis analysis of extant and ancestral PHBases. Lane 1: Molecular weight markers, Lane 2: eI, Lane 3: eII, Lane 4: eIII, Lane 5: eIV, Lane 6: ANC1. B) Primary structure of the ANC1 enzyme. The PDBDase consensus structures are in bold: Oxianion hole – (M/L)xxxHGCxQ, active site histidine – (S/A/G/L)xxxH(P/V/D)xxxG, active site aspartate – GxxDxTV, active site serine – (Q/R)x(I/V)xGLS(A/G)G. X corresponds to any amino acid.

**Figure 4:**
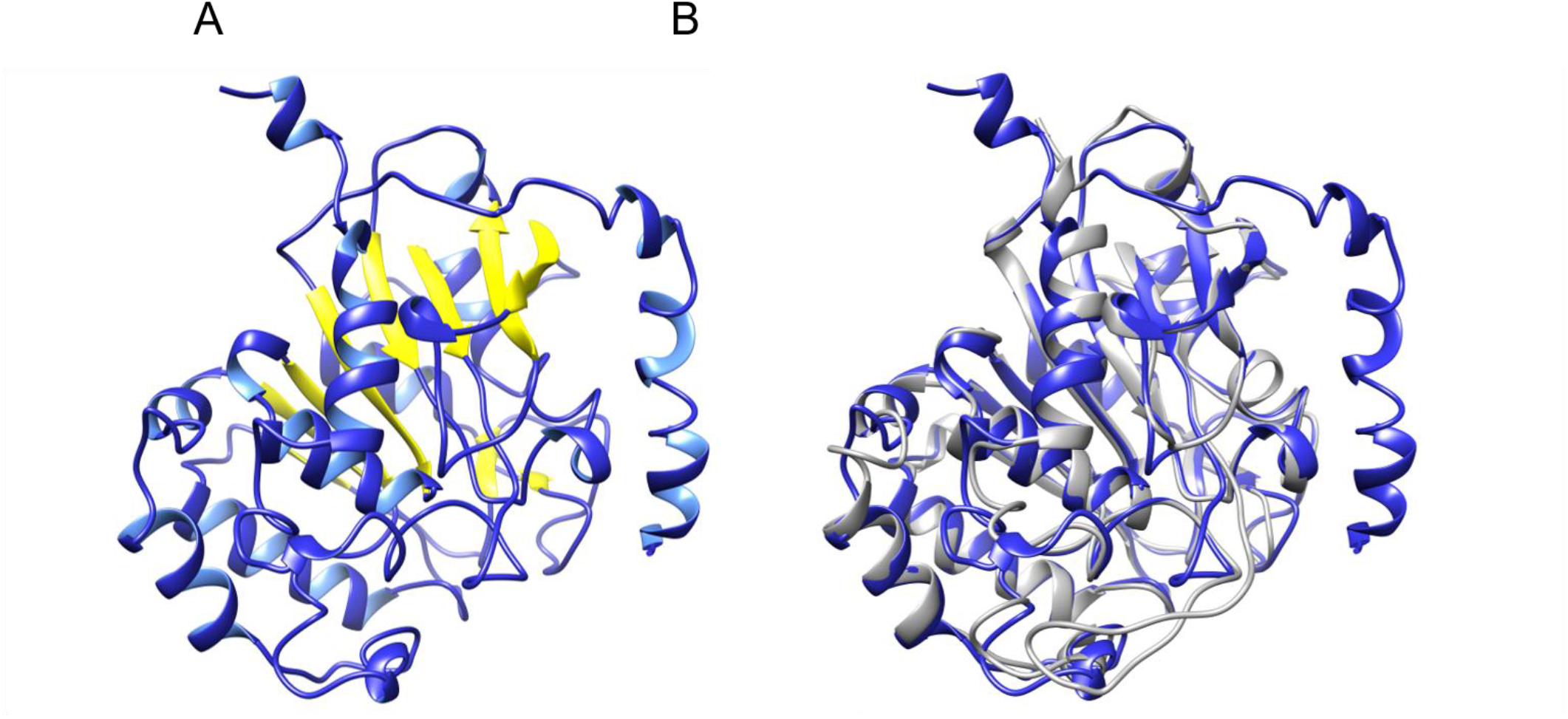
Three dimensional best threaded model of the ANC1 PHBase. Panel A, The NVT relaxed ANC1 model with the central beta strand in yellow. Panel B, 3D superposition of the ANC1 model (blue) and 2D80 (grey).

### 3.3 Enzymatic activity

All of the enzymes are catalytically active against granulated PHB. The turbometric assay measures the loss of light scattering (decrease in absorbance at 650 nm) caused by PHB granules as they are converted into monomeric hydroxybutyrate and small (but soluble) oligomer PHB. As is seen in Figure 5 for ANC1. Comparing the activity parameters of ANC1 relative to the four extant homologs, it is seen that the maximum velocity of the reaction (V_max_) is between 4.4 and 7.3 fold faster and the catalytic rate constant (k_cat_) is between 2.8 and 3.9 fold faster; indicating that the ancestral enzyme is turning over substrate nearly three to four fold faster than are the extant enzymes. Not only is the ancestral sequence kinetically faster, but it also binds to the PHB substrate with between a 1.3 and 1.9 fold lower K_m_. Ancestral sequence reconstruction therefore positively impacts both substrate binding as well as the chemical step in the enzymatic reaction, with a 4.4 to 6.2 fold increase in catalytic efficiency (k_cat_/K_m_). These results are presented in Table III. The thermally induced unfolding reaction is not reversible as all five enzyme homologs precipitate out of solution beyond their temperature maximum. This is most likely due to the existence of disulfide bonds in all of the enzymes. Even though a rigorous thermodynamic analysis in this system is not possible, it is possible to measure enzyme activity as a function of temperature and calculate a temperature of maximum activity. This result is shown in Figure 6. All the extant enzymes have a nearly identical slope during the approach to T_max_, indicating an identical Q_10_ temperature coefficient (in the range between 30 and 40 °C). ANC1 is characterized by a lower slope (and therefore lower Q_10_) in the range of 40 to 50 °C. The ancestral enzyme has a temperature of maximum activity that is between 6.9 and 11.6 degrees Celsius higher than the extant enzymes (Table III).

**Figure 5:**
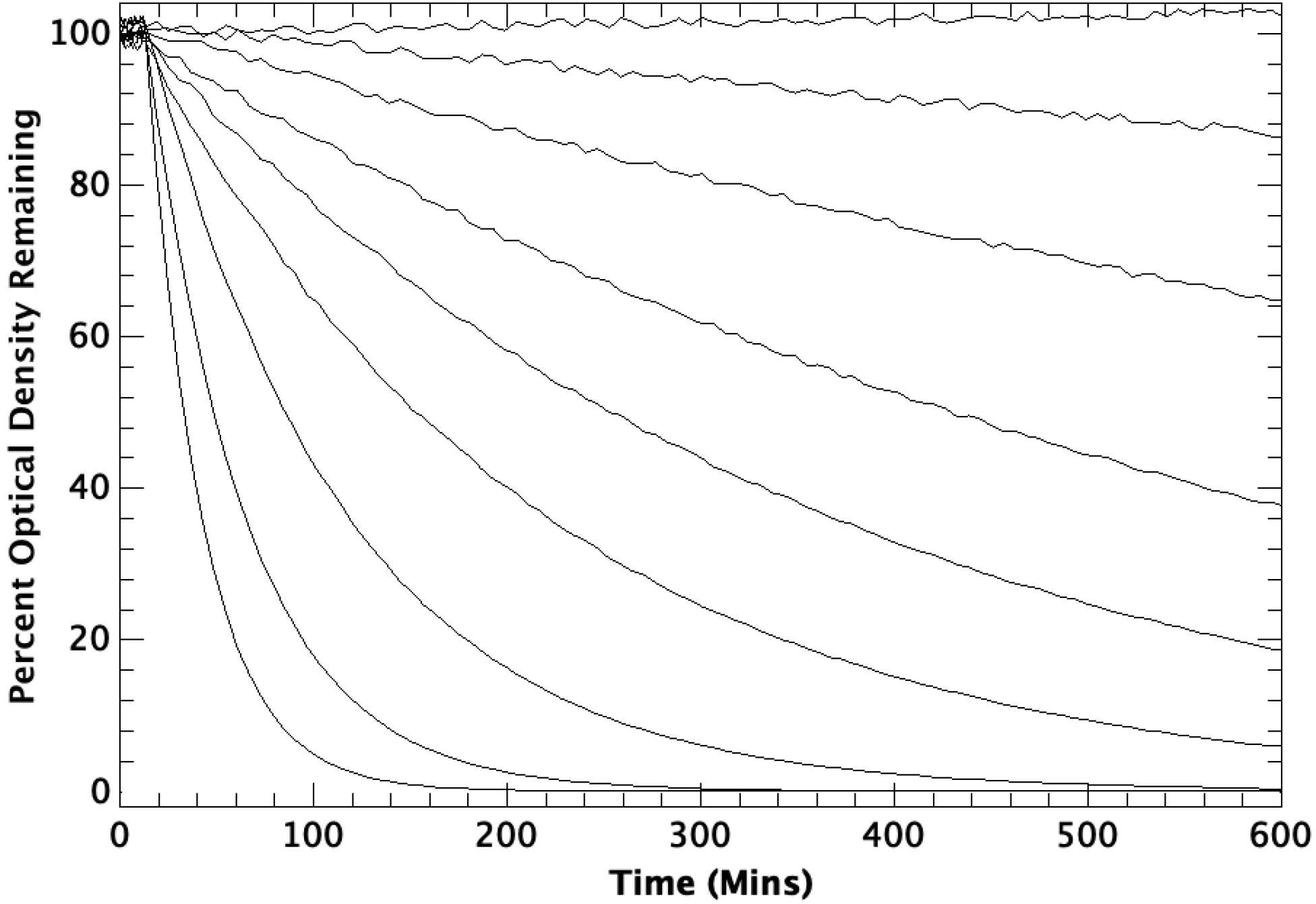
Enzymatic activity as a function of time in the turbidimetric assay. Curves from top to bottom correspond to a total enzyme addition (at t = 10 mins): 0.0 ng, 2.0 ng, 4.0 ng, 6.0 ng, 8.0 ng, 10.0 ng, 12.0 ng, 14.0 and 18.0 ng. Reaction conditions: 25 mM PIPES, pH 6.5, 10 mM CaCl_2_, 300 mM NaCl, 100 mM KCl, 40 °C.

**Figure 6:**
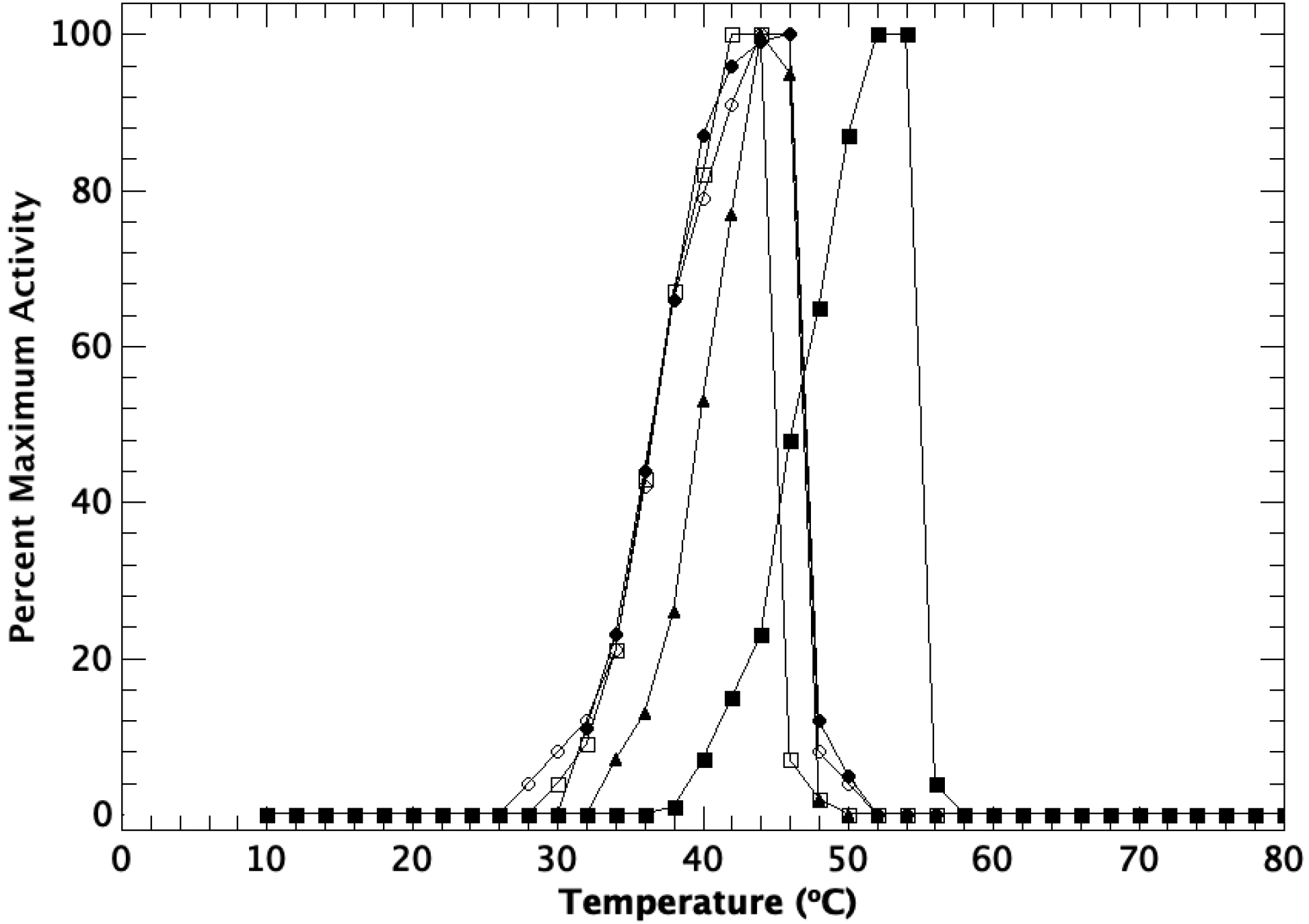
Temperature of maximum activity of extant and ancestral PDBDases. Reaction conditions: 25 mM PIPES, pH 6.5, 10 mM CaCl_2_, 300 mM NaCl, 100 mM KCl Closed circles, eI; open circles, eII; open squares, eIII; triangles, eIV; closed squares, ANC1.

**Table 3.**
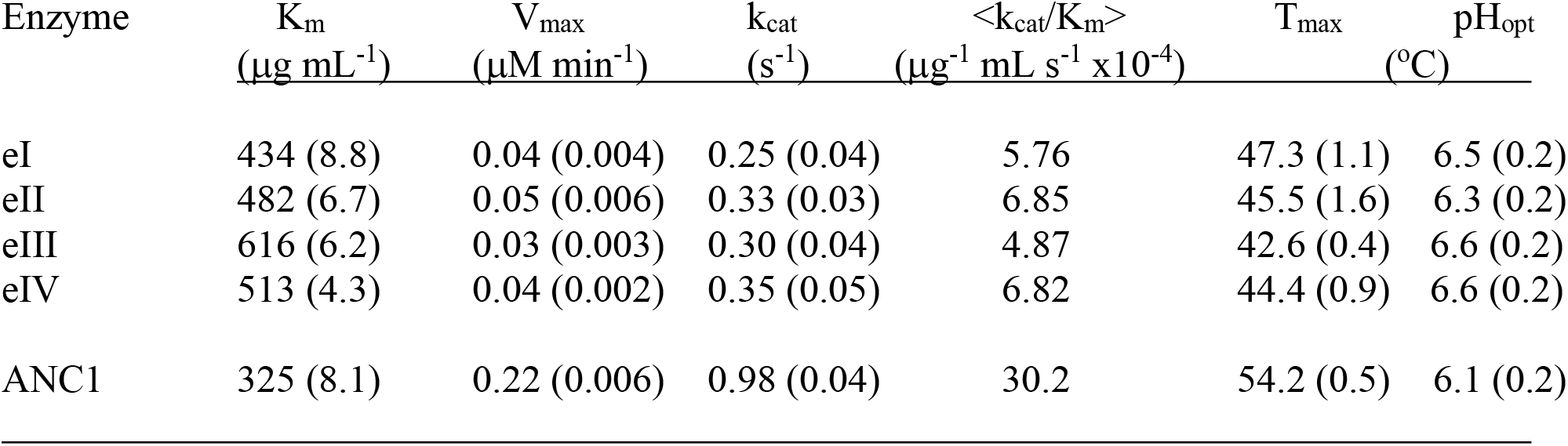
Enzymatic properties of *Halomonas* PHBases

### 3.4 Activity of tethered enzymes

All five PHBase fusion proteins can be coupled to the surface of a Ni-NTA polystyrene bead. However, the enzyme homolog’s ability to depolymerize a PHB film and the enzyme stability over a long course reaction is extremely heterogeneous. Once coupled, eIV shows very little activity against a PHB film, on average 3.3 fold lower activity versus the other extant enzymes (and nearly 10 fold lower than ANC1 activity). The eIV reaction is linear over time. Extant enzymes e1-III show a linear increase in the formation of HB, but all plateau, indicating that the enzymes have stopped working (eI at 36 hours, eII at 56 hours, and eIII at 44 hours). The extant enzymes eI-III are approximately three fold lower in activity compared to ANC1. In contrast ANC1 remains fully active over the 72 hour time course experiment. These results are shown in Figure 7.

**Figure 7:**
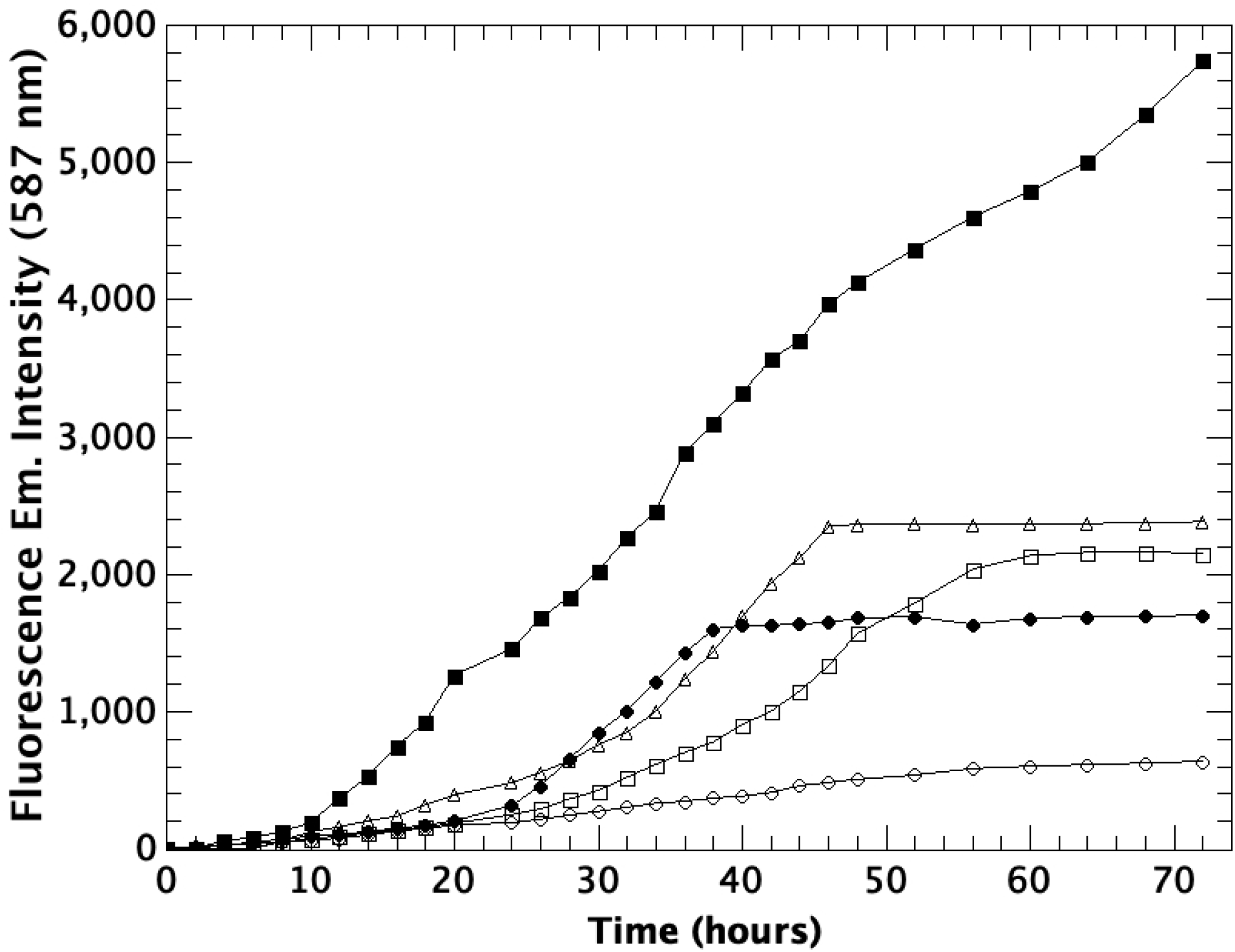
Release of hydroxybutyrate from PHB film using tethered PHBases. Reaction conditions: 25 mM PIPES, pH 6.5, 10 mM CaCl_2_, 300 mM NaCl, 100 mM KCl, 40 °C. Closed circles, eI; open circles, eII; open squares, eIII; triangles, eIV; closed squares, ANC1.

## 4. Discussion

Evolutionary analysis [55]when used in conjunction with ancestral sequence reconstruction [56] offers an alternative technique to site directed mutagenesis or directed evolution [57]. It can also be used to identify an enzyme candidate that is then further modified using site directed mutagenesis. This is important as many naturally occurring enzymes are not directly suited for industrial applications due to low catalytic efficiency and/or stability in industrial environments [58]. A hallmark of ASR is in identifying proteins with increased thermostability, which is a goal for many industrial enzyme applications. Presumably this observation is due to the substantially hotter Earth in earlier eras [59–61], but there is some disagreement on the origin of ASR increased thermostability [62] as it relates to both evolutionary selection pressure and the how phylogenies are calculated. However, it has been observed that when ASR is applied to more ancient evolutionary nodes (*e.g.-* Precambrian) the T_m_s for these proteins are 30 –35 °C higher than their extant counterparts [59,60]. This is too large of an increase to be accounted for simply based on ancestral sequence reconstruction methods and any inherent biases [63]. In some systems it is the extant enzyme that is characterized by a greater thermostability than the ASR sequence [61]. An increase in thermostability can be driven by better core packing, decreased loop mobility, surface charge reorganization, disulfide bonding, and/or changes to the folding pathway trajectory [64,65]. In the case of extant versus ancestral PHBases, the driving mechanism is not fully apparent. In any case, the technique offers insights to enzyme plasticity [66]. All enzymes have a number of disulfides and a similar isoelectric point. So, in this system some other stabilizing factor may predominate. In the past it was thought that thermal stability was inversely related to catalytic ability due to a loss of flexibility associated with higher stability. However, Roca *et al*. [67] have shown that the rate of the chemical step is a function of the activation barrier and the amount of reorganization energy and is not related to the degree of flexibility. In fact, they also showed that the more catalytically efficient enzymes display less flexibility/motion during catalysis. The role of structural dynamics in catalysis remains an open question [68].

We have not attempted to determine unfolding pathways or perform structural studies. Nonetheless, the method is extremely promising to rapidly identify candidate enzymes for industrial processes. It is possible to utilize information derived from the ASR calculation to predict additional mutations that drive increased thermostability [32]. This method affords a more data-driven approach to site directed mutagenesis of the identified candidate enzyme than random mutagenesis.

The evolution of catalytic efficiency and enzymatic velocity / substrate turnover are harder questions to resolve directly from ASR. There are many factors that contribute to the ability of an enzyme to bind a substrate, perform a chemical step, and release product [69]. These pleiotrophic contributors are harder to identify in a phylogenetic and ASR analysis than is thermostability. The diversity of active site architectures, even within an enzyme class, indicates that chemo-physiological pressures, driven in part by environments and evolutionary constraints alter catalytic performance to match metabolic demands [70,71]. Hence there is not a driving force like the cooling Earth to inversely drive thermostability [72], rather catalytic descriptors are probably stochastically arranged throughout evolutionary history. Catalysis is primarily driven by transition state theory (for a review see[73]), which in turn is dependent on active site geometry and composition, binding affinity, and the energetics along the reaction coordinate. Directional drives in catalytic rates across a phylogeny are also made more difficult to identify due to the role that enzyme promiscuity plays in the evolution of enzyme structure and function [74], yet clearly it is possible to understand the evolution of catalytic function through phylogenetic sequence [75] and evolutionary analysis [76]. A functional strategy for the use of ASR to identify candidates for industrial processes may be to first identify a region of evolutionary space to begin the analysis and focus on identifying a candidate ancestral enzyme with a measured stability in a range of interest, followed by focusing on optimizing enzymatic activity for the process at hand.

Therefore, the starting point for ASR is an important consideration. There are 6,415 PHB depolymerase sequences in the NCBI Identical Protein Group database. It is certainly possible to perform a phylogenetic analysis on all of them simultaneously, but such a large dataset offers a diluted return for the computational investment. A preferred approach, as performed here, is to pick a smaller sequence space subset starting with a family, group of genera, or a single genus of interest. Having knowledge of the ultimate use, catalytic and stability requirements, and the industrial environment in which the enzyme will function can help narrow the choice. Some initial search areas would be extremophiles [77,78]; including organisms that live at high temperatures, at low temperatures, in alkaline or acidic environments, or in high salinity conditions. Often the Archaea are a promising first analysis area (*e.g.-* [79]), particularly the Haloarchaea [80]. Choice of starting point is more critical when the final bioreactor process will utilize an intact organism versus a purified enzyme. The size of the initial database is also dependent on computational resources and the number of candidate enzymes that can be constructed and screened. We have found in practice that keeping the number of species to below ~200 results in striking a balance between minimizing the total number of nodes and having enough evolutionary diversity to reconstruct meaningful (and useful) ancestor sequences.

## 5. Conclusions

All the parameters, K_m_, k_cat_, T_m_, solubility, charge, that are subject to optimization for any industrial process can be addressed by the analysis of ancestral sequence – regardless of the mechanism that created them or their evolutionary validity. In the present study, an ancestral sequence was reconstructed that displayed a lower K_m_, a higher catalytic turnover, and an increased maximum temperature of activity than an extant enzyme from the same genus. This single sequence has improved upon the parameters that need to be optimized for a large scale industrial enzymatic process. The ANC1 sequence is in fact a prime candidate for a bioreactor process for the depolymerization of PHB plastics.

## Supporting information

All supplemental data

## Contributors

SQ: Research design, phylogenetic analysis, enzyme preparation and characterization, and writing. KK: Python script development and testing, research design, and writing.

## Acknowledgements

This work was solely funded by the Kimberly-Clark Corp.

