## Supplementary material for "The use of phylogenetics and ancestral sequence reconstruction to identify improved halophilic enzymes for the biodegradation of poly(*R*-3-hydroxybutyrate)": All supplemental data

Python script for sequence population analysis:

```
from Bio import SeqIO
from Bio.SeqRecord import SeqRecord
from Bio.SeqUtils import ProtParam
from Bio.SeqUtils import IsoelectricPoint

def output_proteinDem(filename):
    x={}
    for record in SeqIO.parse(filename, "fasta"):
        Cysteine=0
        Neg=0
        Pos=0
        Aromatic=0
        seq1=str(record.seq)
        X = ProtParam.ProteinAnalysis(seq1)
        iePt=X.isoelectric_point()
        length=len(seq1)
        for i in seq1:
            if i=='C':
                Cysteine=Cysteine+1
            if i=='D' or i=='E':
                Neg=Neg+1
            if i=='K' or i=='R':
                Pos=Pos+1
            if i=='W' or i=='F' or i=='Y':
                Aromatic=Aromatic+1

x[record.description]=iePt,length,Cysteine,Neg,Pos,Aromatic
            with open('output_proteinDemographics.txt', 'w') as f:
                f.writelines('Enzyme\tIsolelectric point\tNo. of
AAs\tCysteine\tNegative AAs\tPositive AAs\tAromatic AAs\n')

f.writelines('{}\t{}\t{}\t{}\t{}\t{}\t{}\n'.format(k,v[0],v[1],v[2],v
[3],v[4],v[5]) for k,v in x.items())

output_proteinDem("halomon.fasta")
```

Python script for truncating sequence names:

```
from Bio import SeqIO
from Bio.SeqRecord import SeqRecord

original_file = "halomon.fasta"
corrected_file = "halomon-short.fasta"

with open(original_file) as original, open(corrected_file, 'w') as
corrected:
    records = SeqIO.parse(original_file, 'fasta')
    for record in records:
        record.description=record.id
        #print(record.description)
        SeqIO.write(record, corrected, 'fasta')
```

Python script for selecting particular files out of the master database:

```
import numpy as np #read in the names of the enzymes from file
2enzymes = np.genfromtxt('The-1057-Names.prn', dtype=None,
usecols=(0,))enzymes = [a.decode("utf-8") for a in enzymes] #format
the string in a better way#open the total data file to read line by
line and create a new filewith open('all.fasta', 'r') as file,
open('selected.fasta', 'w') as new_file: #change name of
selected.fasta if wanted for line in file: #read line by line
     for enzyme in enzymes: #loop over all the enzymes in the list
          if line.startswith('>{}'.format(enzyme)): #find the entry
for the enzyme                new_file.write('{}'.format(line))
#write line to new file                #now copy the next lines so
that you get the whole entry                next_line = next(file)
               while '>' not in next_line: #stop copying lines when
we get to the next entry
     new_file.write('{}'.format(next_line))
     next_line = next(file)
```

CLUSTAL multiple sequence alignment by MUSCLE (3.8)

```
WP_089674669.1      ------------MGIAVHQRRLMAALILLGSAVAHAE-----EEAPGLPALGAANDQASV
WP_041159480.1      ---------------MPTSSNSHDRLTNLQ------------------------------
WP_083700770.1      MIDA---QRMQEATRLTRAGQLHEAMAILQGAMPDGKSTGSSEHAPDASYSAGNTFTGTC
WP_035563078.1      MIDATMTKRMEEATRLTRAGKLQEAMAVLQGSAPESS----GVETPNTSYQEGNTFDGTY
ANC1                MIDATMTKRMEEATRLTRAGKLHEAMALLQGAMPEAK----GEETPDNPYQGGNTFEGTC
                                        .    :  *

WP_089674669.1      VGVS--------------------------------SGGYMASQLAVAWPERFSGV----
WP_041159480.1      ------------------------------------------------------------
WP_083700770.1      EVVDEEVSSASAEGHSTAEPSAGAAPFTAFGHARQATAGTAGGGMPSAWRDTWRQMRESM
WP_035563078.1      EVIDEEASESKPE----P----AEAPAGAF------LPGAI--KVPAAWRDKWMKFRGGM
ANC1                QVVDEEVTASKPK----PEPESAEATSGAF------SSGDI--KMPAAWRDKWMKFRGAM


WP_089674669.1      ------------------GMLAAGPWGCAQGALSLALN-QCMMTRRGLPSLDELEQRRER
WP_041159480.1      ------------------------QFGADPGSLHADTYIPKNFPKNG-PLVVVLHGSTQS
WP_083700770.1      ARSELS-----TSEIAQPGAFTAGRFSHPAGTREYKLYLPGGYHGQALPLVVMLHGCTQD
WP_035563078.1      AQPEQAPN---TSEEPQPGAFTAGRFTNPVGSRDYKLYIPSGYTGQALPLVVMLHGCTQN
ANC1                AQPEQAPSEFGTTEEPQPGAFTAGRFSNHVGTRDYKLYIPSGYHGQALPLVVMLHGCTQN
                                         :    *:             .. * :  *    :

WP_089674669.1      YLSLDQVGSQDALSQLRAF-VWHGDADETVSPALGDLLAQQWQGWLESPEQQLRYVQRAN
WP_041159480.1      AEGYDRGSGWSTLADESGIALLYPKQRKTNNP------ISSFN-WFKSGDSR--------
WP_083700770.1      PDDFAAGTRMNRLAEEQQFCVLYPAQPMTANS------SKCWN-WFKAEDQQ--------
WP_035563078.1      PDDFAAGTEMNRLAEEQLCYVLYPAQPMTANS------SKCWN-WFKAEDQQ--------
ANC1                PDDFAAGTNMNRLAEEQLFCVLYPAQPMTANS------SKCWN-WFKAEDQQ--------
                      .       . *::     : :     * ..       . :: *::: :..

WP_089674669.1      TGHGWPVAMPKDAPIDPQSLGDCRNGGGSHVLACGEDVAGEM---MAWLYPE--RETNAS
WP_041159480.1      RGGGEPLSIRHMI----KQVVDDHAIDSTRVFVTGMSSGGAMTSVMLATYPEVFAGGAII
WP_083700770.1      RGGGEPAILAGMT----RQIIDTHALDASRVYVAGLSAGGAMATTLAMTYPDLYAAVGVH
WP_035563078.1      REGGEPAIIAGMT----RQIIDTHGLDGTRIYVAGLSAGAAMATTLAMTYPDLFAAVGVH
ANC1                REGGEPAILAGMT----RQIIDTHGLDASRVYVAGLSAGAAMATTLAMTYPDLFAAVGVH
                       * *  :        ..: * .  ..:.: . * . .. *   :   **:

WP_089674669.1      EGELLAFDQSDFAAKGFADTGYVFVPEACEA---GGC------PVTVALHG---CQMNAE
WP_041159480.1      AGLPYRSADNLIEAMVRMKGYGGPSDHRLDALVRGASTFDGPWPTISVWHGGSDSTVDNA
WP_083700770.1      SGLPHAVAQSLPDALAAMQGGR-PMAAASSA---GASTWASTVPAV-VFHGDRDTTVHPS
WP_035563078.1      SGLPHGVAQSLPDALGAMQGGTGPL-GGGKA---KGAGWASEVPAI-IFHGDLDTTVHPS
ANC1                SGLPHGVAKSLPDALGAMQGGTGPLGGGGKA---QGAGWASEVPAI-IFHGDRDTTVHPS
                     *      ..   *    .          .*    ..      *.    **     :

WP_089674669.1      AIDDTFVRYSGLNRWAA--EHGQVVL---YPQAESSMANPQACWDWW-----GFAESTW-
WP_041159480.1      NADSIVRQWQQIHKVEGPPTRVEQVAG--FPRQVWCNADRQEVIEEYIIEGMGHGTPIMA
WP_083700770.1      NADRVAAQYAVSRGAGTSLERGQVANGHAYTKTLHHDAHGQLCLEQWVIHGAGHA---WS
WP_035563078.1      NADRVAAQYSTSRQESATVEQGKVANGHAYTRTTHHNEKGEPHMEQWRIHGAGHA---WS
```

```
ANC1                NADRVAAQYAASRQARVTVEQGKVANGHAYTRTTHHDAKGKPRLEQWRIHGAGHA---WS
                      *    .:   .       . : .    :..        :   : :     *..

WP_089674669.1      -------QINPLHDTRDGTQTQALMAM--LDH-LQSATANKAATAE
WP_041159480.1      GGDEGLGEADKYMLEVGISSTRHIAHFWGLTE--------------
WP_083700770.1      GGD----ARGSHTDAKGPDATQEMLRFF-QSHALSVDQAGRSAT--
WP_035563078.1      GGS----AKGSYTDSKGPDATKEMLRFF-LQH----QQCERDMN--
ANC1                GGS----AKGSYTDPKGPDATKEMLRFF-LQH----QQAERSASSS
                             .      .   *. :  :
```

Bootstrapped ML tree (1000 replicates) showing percentage of trees with the same node:

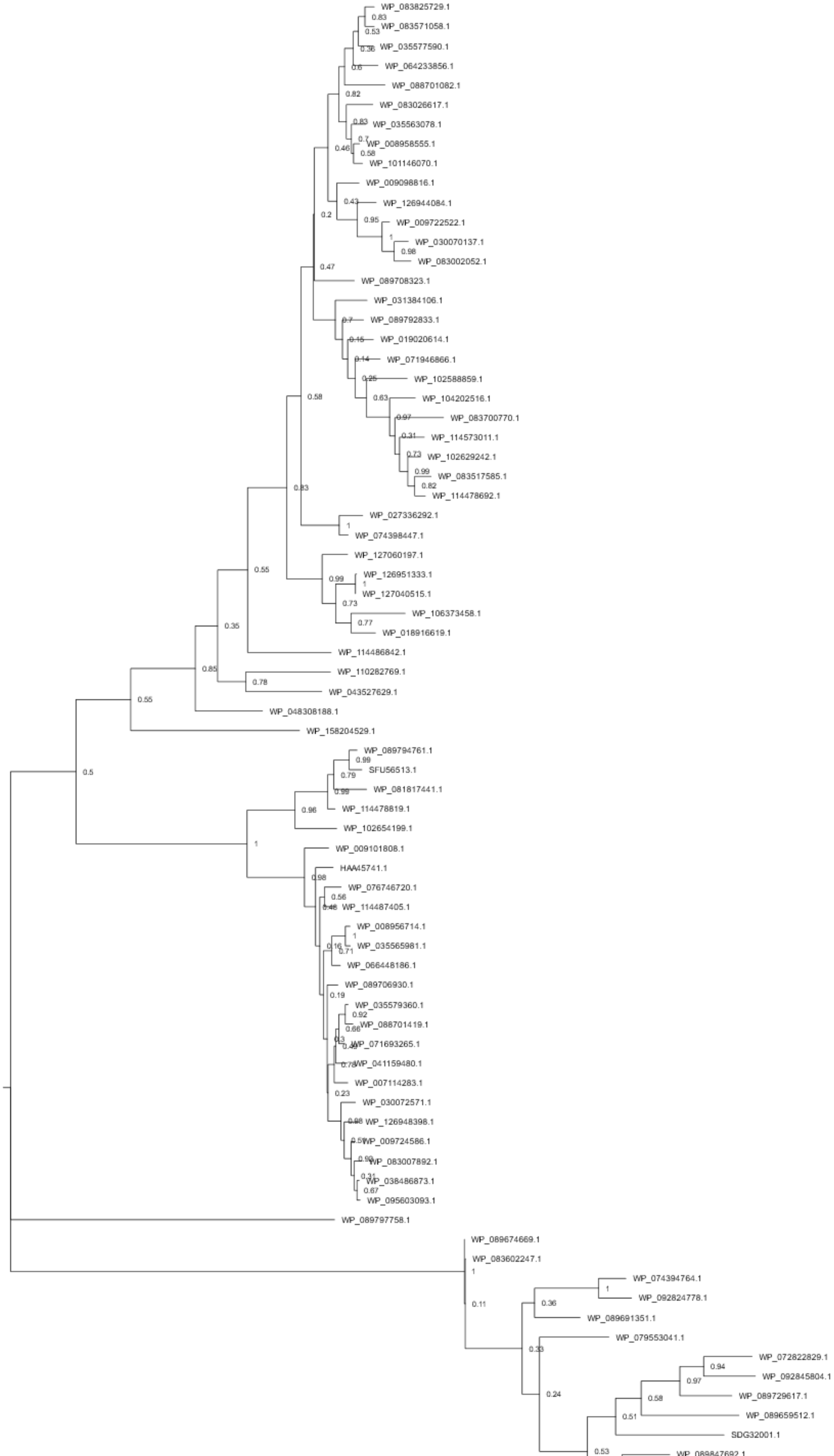
WP_083825729.1
WP_083571058.1
WP_035577590.1
WP_064233856.1
WP_088701082.1
WP_083026617.1
WP_035563078.1
WP_008958555.1
WP_101146070.1
WP_009098816.1
WP_126944084.1
WP_009722522.1
WP_030070137.1
WP_083002052.1
WP_089708323.1
WP_031384106.1
WP_089792833.1
WP_019020614.1
WP_071946866.1
WP_102588859.1
WP_104202516.1
WP_083700770.1
WP_114573011.1
WP_102629242.1
WP_083517585.1
WP_114478692.1
WP_027336292.1
WP_074398447.1
WP_127060197.1
WP_126951333.1
WP_127040515.1
WP_106373458.1
WP_018916619.1
WP_114486842.1
WP_110282769.1
WP_043527629.1
WP_048308188.1
WP_158204529.1
WP_089794761.1
SFU56513.1
WP_081817441.1
WP_114478819.1
WP_102654199.1
WP_009101808.1
HAA45741.1
WP_076746720.1
WP_114487405.1
WP_008956714.1
WP_035565981.1
WP_066448186.1
WP_089706930.1
WP_035579360.1
WP_088701419.1
WP_071693265.1
WP_041159480.1
WP_007114283.1
WP_030072571.1
WP_126948398.1
WP_009724586.1
WP_083007892.1
WP_038486873.1
WP_095603093.1
WP_089797758.1
WP_089674669.1
WP_083602247.1
WP_074394764.1
WP_092824778.1
WP_089691351.1
WP_079553041.1
WP_072822829.1
WP_092845804.1
WP_089729617.1
WP_089659512.1
SDG32001.1
WP_089847692.1
